# Persistent inflammation promotes endocannabinoid release and presynaptic cannabinoid 1 receptor desensitization

**DOI:** 10.1101/2022.06.20.496891

**Authors:** Courtney A. Bouchet, Aaron Janowsky, Susan L. Ingram

## Abstract

Pain therapies targeting the cannabinoid system are increasing with expansion of cannabis legalization but adaptations in the endogenous cannabinoid system during inflammatory pain could limit their efficacy. Presynaptic inhibition of GABA release mediated by cannabinoid 1 receptor (CB1R) agonists in the ventrolateral periaqueductal gray (vlPAG) is markedly reduced in male and female Sprague Dawley rats after persistent inflammation induced by Complete Freund’s Adjuvant (CFA). Inflammation results in increased endocannabinoid (eCB) synthesis and desensitization of presynaptic CB1Rs that is reversed by a GRK2/3 inhibitor, Compound 101. Despite CB1R desensitization, eCB activation of CB1Rs is maintained after inflammation. Depolarization-induced suppression of inhibition (DSI) in naïve animals is rapid and transient, but is prolonged in recordings after inflammation. Prolonged DSI is mediated by 2-arachidonoylglycerol (2-AG) indicating reduced monoacylglycerol lipase (MAGL) activity. These adaptations within the endogenous cannabinoid system have important implications for the development of future pain therapies targeting CB1Rs or MAGL.

## Introduction

The cannabinoid 1 receptor (CB1R) is one of the most highly expressed GPCRs in the brain (Herkenham et al 1990) and is primarily localized to presynaptic terminals, where its activation inhibits neurotransmitter release (Katona et al 1999, Mikasova et al 2008, Vaughan & Christie 2005, Vaughan et al 2000). CB1Rs are activated by endogenous cannabinoid ligands, endocannabinoids (eCBs) that negatively regulate synaptic transmission through on-demand synthesis, retrograde transport and activation of CB1Rs. eCB activation of CB1Rs is tightly controlled by enzymes responsible for eCB on-demand synthesis and rapid degradation (for review see Ahn et al 2008). The two most well studied eCBs are 2-arachidonylglycerol (2-AG) and anandamide (AEA).

In the brain, levels of 2-AG are more than 100 times higher than AEA (Stella et al 1997). 2-AG is regulated by the synthesis enzyme, diacylglycerol (DAGL; Bisogno et al 2003) and the catabolism enzyme, monoacylglycerol lipase (MAGL; Dinh et al 2002, Dinh et al 2004). Through this endogenous cannabinoid system, eCBs and the CB1R regulate neurotransmitter release from the presynaptic terminal.

Expression of eCBs and their degradation enzymes are altered by inflammation in several brain areas (Vecchiarelli et al 2021). Our prior study demonstrated a reduction in CB1R suppression of GABA release that was the result of reduced protein expression in the rostral ventromedial medulla (RVM) with persistent inflammation (Li et al 2017). The RVM is integral to descending pain modulation and, along with the ventrolateral periaqueductal gray (vlPAG), constitutes the descending pain modulatory pathway. Within the vlPAG, CB1Rs are densely expressed (Wilson-Poe et al 2012) and their activation modulates neurotransmitter release in naïve animals (Aubrey et al 2017, Drew et al 2009, Lau et al 2014, Vaughan et al 2000, Wilson-Poe et al 2015), but adaptations in the cannabinoid system after persistent inflammation in the vlPAG are not understood. Therefore, we sought to investigate how persistent hindpaw inflammation impacts cannabinoid regulation of GABA release within the vlPAG.

The present results describe an inflammation-induced increase in eCB levels in the vlPAG, leading to desensitization of CB1Rs by a GRK2/3-dependent mechanism. While this desensitization is clearly observed with exogenous agonists, endogenous release of 2-AG continues to induce CB1R-dependent suppression of inhibition after inflammation. Compared to naïve, the eCB-induced suppression is prolonged after persistent inflammation. Together, results show a distinction between CB1R activation by exogenous and endogenous cannabinoid ligands, as well as alteration in the endogenous cannabinoid system in the vlPAG after persistent inflammation. These adaptations have important implications for future therapeutic drug development.

## Results

### Persistent inflammation reduces CB1R inhibition induced by exogenous agonists

Plasticity within the cannabinoid system induced by persistent inflammation in the vlPAG was examined following Complete Freund’s Adjuvant (CFA) injection into the hindpaw of male and female Sprague Dawley rats. All experiments were conducted 5- 7d after CFA injection. Whole-cell patch clamp recordings of electrically evoked inhibitory postsynaptic currents (eIPSCs) were used to measure GABA IPSCs and the inhibition of GABA release by the non-selective cannabinoid receptor agonist, WIN- 55,212-2 (WIN; 3 μM). In tissue from naïve animals, WIN reduced eIPSC amplitudes by 57 ± 5% compared to baseline (Fig. 1A,B). CFA-induced inflammation significantly reduced WIN-mediated inhibition of eIPSCs to 18 ± 4% (Fig. 1A,B). WIN inhibition was reversed by the CB1R selective antagonist, SR141716A (rimonabant, RIM; 3 μM). No sex differences were observed in WIN-mediated suppression of GABA release in recordings from either naïve or CFA-treated rats (Fig. S1), so data from male and female rats were combined for all analyses. There were no differences in baseline eIPSC paired pulse ratios (unpaired t-test, *t*_13_=0.59; p=0.6) or decay kinetics (unpaired *t*- test, *t*_11_ = 1.0; p=0.3) between recordings from naïve and CFA-treated animals.

**Figure 1:**
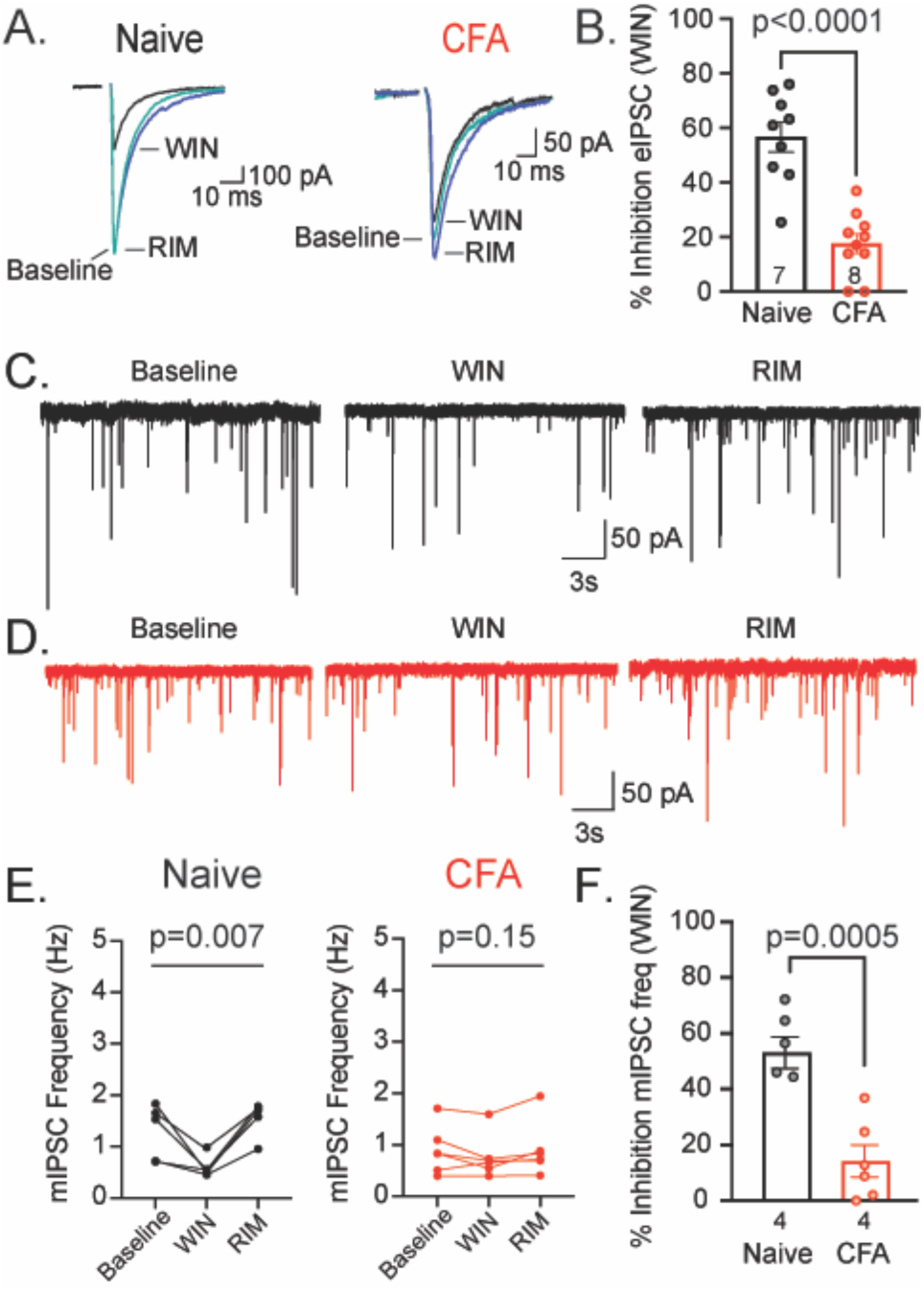
Persistent inflammation reduces WIN-induced inhibition of GABA release. (A) Representative traces of eIPSCs isolated in NBQX recorded from vlPAG neurons in baseline (5μM; teal), the cannabinoid receptor agonist WIN,55212 (WIN 3 μM; black) and the CB1R selective antagonist rimonabant (RIM; 3 μM; blue) from naïve (left) and CFA-treated (right) animals. (B) Percent inhibition of eIPSC amplitude by WIN in slices from naïve (black bar) and CFA-treated rats (red bar) (unpaired t-test, *t*_14_=5.34; p=0.0002). (C,D) Representative trace of mIPSCs recorded from vlPAG neurons in baseline containing TTX (500 nM) and NBQX (5 μM), WIN (3 μM), and RIM (3 μM) from slices of naïve (black, C) or CFA-treated rats (red, D). (E) mIPSC frequency at baseline, WIN, and RIM from slices of naïve (black) and CFA-treated (red) rats. (F) WIN percent inhibition of mIPSC frequency from naïve (black) and CFA-treated (red) rats (unpaired t-test, t_(10)_=4.65; p=0.0009).

To determine whether inflammation also affects spontaneous GABA release and the inhibition of spontaneous release by CB1Rs, we measured miniature IPSCs (mIPSCs) in the presence of tetrodotoxin (TTX; 500 nM). WIN suppressed mIPSC frequency by 56 ± 5% in tissue from naïve animals and this effect was significantly reduced (14 ± 6%) after persistent inflammation (Fig. 1C-F). Activating CB1Rs had no effect on mIPSC amplitude (One-way ANOVA: F(1.1, 5.5)=0.43; *p*=0.56), consistent with a presynaptic effect of CB1R activation.

To determine whether the reduction in CB1R suppression of GABA release is due to a general change in presynaptic GPCR signaling or downstream signaling pathways, we investigated the effects of persistent inflammation on the cannabinoid 2 receptor (CB2R) and presynaptic μ-opioid receptor (MOR) inhibition of GABA release. The CB2R agonist, AM1241 (3 μM) did not affect mIPSC frequency in vlPAG slices from naïve animals (14 ± 4% inhibition; Fig. 2A) and this was not changed after persistent inflammation (17 ± 10% inhibition; Figure 2A,B; unpaired t-test, *t*_15_=0.71; *p*=0.5). While CB2R activation does not affect GABA release within the vlPAG, MOR activation suppresses GABA release to a similar extent in both naïve and CFA-treated slices. The MOR selective agonist DAMGO (1 μM) inhibited eIPSCs to the same extent in slices from naïve and CFA-treated rats (Naïve: 69 ± 16%; CFA: 66 ± 23%; Fig. 2C,D).

**Figure 2:**
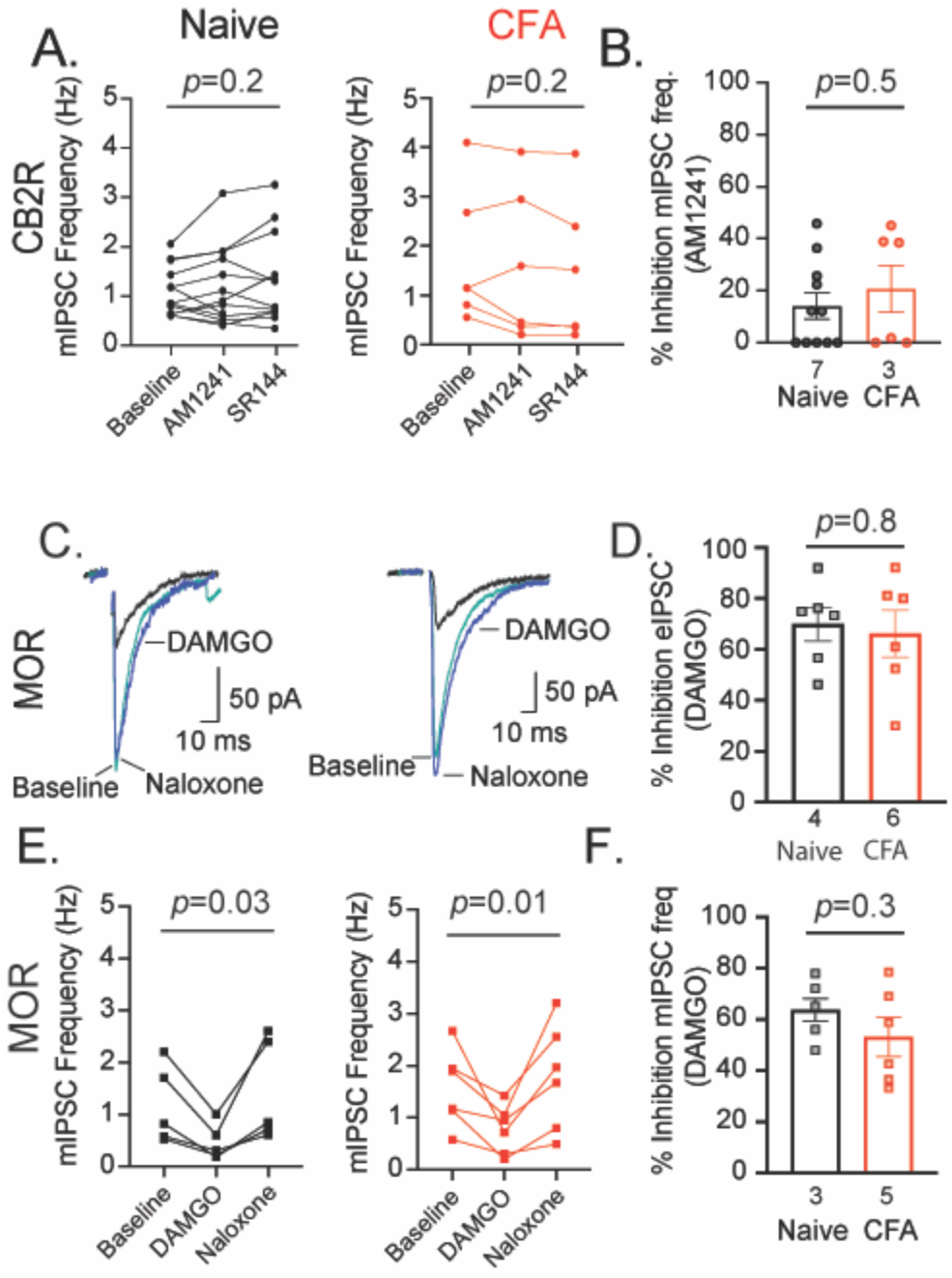
Persistent inflammation does not affect CB2R or MOR suppression of GABA release. (A) spontaneous mIPSC frequency in slices from naïve (black) or CFA-treated (red) animals during baseline, CB2R agonist AM1241 (3 μM) and CB2R antagonist SR144528 (3 μM). (B) mIPSC frequency inhibition by AM1241 (unpaired t-test; *t*_15_=0.71; *p*=0.5). (C) Representative eIPSC traces at baseline (5 μM; teal), DAMGO (1 μM; black) and naloxone (1μM; blue). (D) Percent inhibition of eIPSCs by DAMGO in naïve (black bar) and CFA-treated (red bar) conditions (unpaired t-test, *t*_10_= 0.32; *p*=0.8). (E) spontaneous mIPSC frequency in slices from naïve (black) or CFA-treated (red) animals during baseline, DAMGO (1μM), and naloxone (1μM). (F) mIPSC frequency inhibition by DAMGO (unpaired t-test, *t*_9_=1.11; *p*=0.3). Error bars represent SEM, dots indicate individual recordings and numbers represent the number of rats represented per bar.

DAMGO-induced suppression of mIPSC frequency was also unaffected by persistent inflammation (naïve: 64 ± 12%; CFA: 53 ± 18%; Fig. 2E,F). Together, these data indicate that the effects of persistent inflammation are selective to the CB1R within the vlPAG.

### Cannabinoid receptor expression is unchanged following persistent inflammation

Persistent inflammation downregulates CB1R protein in the RVM (Li et al 2017), so we hypothesized that persistent inflammation downregulates CB1R expression in the vlPAG as well. Expression levels were measured using radioligand binding with [^3^H]CP- 55,940. Since this is a different ligand than previously used, we first replicated our findings from Fig. 1 and found that CP-55,940 suppression of GABA release is significantly reduced after persistent inflammation (Naïve: 50 ± 5%, CFA: 4 ± 4%; Fig. 3A,B). Radioligand binding was then carried out using [^3^H]CP-55,940 in vlPAG dissected from naïve and CFA-treated. Surprisingly, there was no difference in total cannabinoid receptor binding (Fig. 3C; Naïve B_max_: 785 ± 61 fmol/mg; CFA B_max:_ 708 ± 126 fmol/mg) or the dissociation constant (Naïve K_d_ = 1.8 ±0.3 nmol; CFA K_d_ = 1.7 ± 0.4 nmol) in vlPAG tissue from naïve and CFA-treated animals. Similarly, persistent inflammation did not impact cannabinoid receptor binding in the dorsolateral striatum or hypothalamus (Fig. S2).

**Figure 3:**
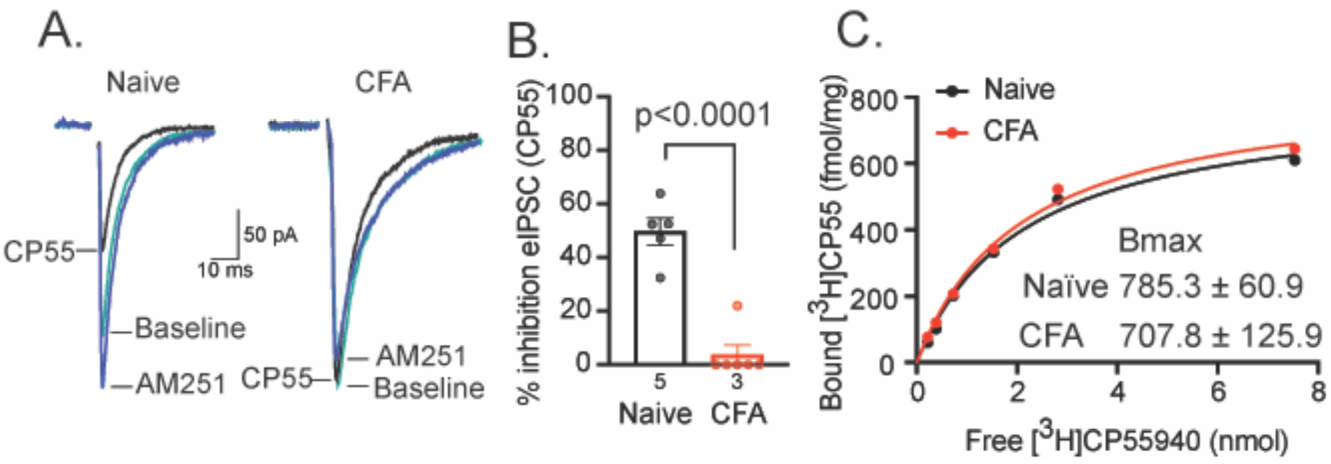
Cannabinoid receptor binding is unaffected by persistent inflammation. (A) Representative traces of eIPSC recorded from vlPAG neurons in baseline containing NBQX (5μM; teal; Baseline), cannabinoid agoinst CP-55,940 (3μM; black; CP55), and CB1-selective antagonist AM251 (3μM; blue) from naïve and CFA-treated rats. (B) Percent inhibition of eIPSC amplitude by CP-55,940 in vlPAG slices from naïve (black bar) or CFA-treated (red bar) rats (unpaired t-test, *t*_9_=7.8; p<0.0001). (C) Representative radioligand binding saturation curve with [^3^H]CP-55,940 and vlPAG tissue from naïve (black) and CFA-treated (red) rats (vlPAG from 8 rats pooled per curve, statistics run on average of 3 curves). Error bars represent SEM, dots indicate individual recordings and numbers represent the number of rats per bar.

### CB1Rs do not display acute desensitization to exogenous agonist

The observation that total cannabinoid receptor binding was unchanged in slices from CFA-treated rats suggested that CB1Rs might be desensitized with persistent inflammation. Similar to other presynaptic GPCRs, CB1Rs do not desensitize during 30 minutes of WIN (3 μM; Fig. 4A). To test whether CB1Rs in the vlPAG desensitize with multiple hours of agonist exposure, slices containing vlPAG were incubated in WIN (3 μM) for 90 minutes up to 5.5 hours and RIM was used to determine the extent of inhibition by WIN over time. RIM increased eIPSC amplitudes similarly after 15 minutes of WIN (275 ± 48%) or >1 hour of WIN (274 ± 92%; Fig. 4B,C). These results indicate that CB1Rs are resistant to desensitization, even after several hours of WIN exposure.

**Figure 4:**
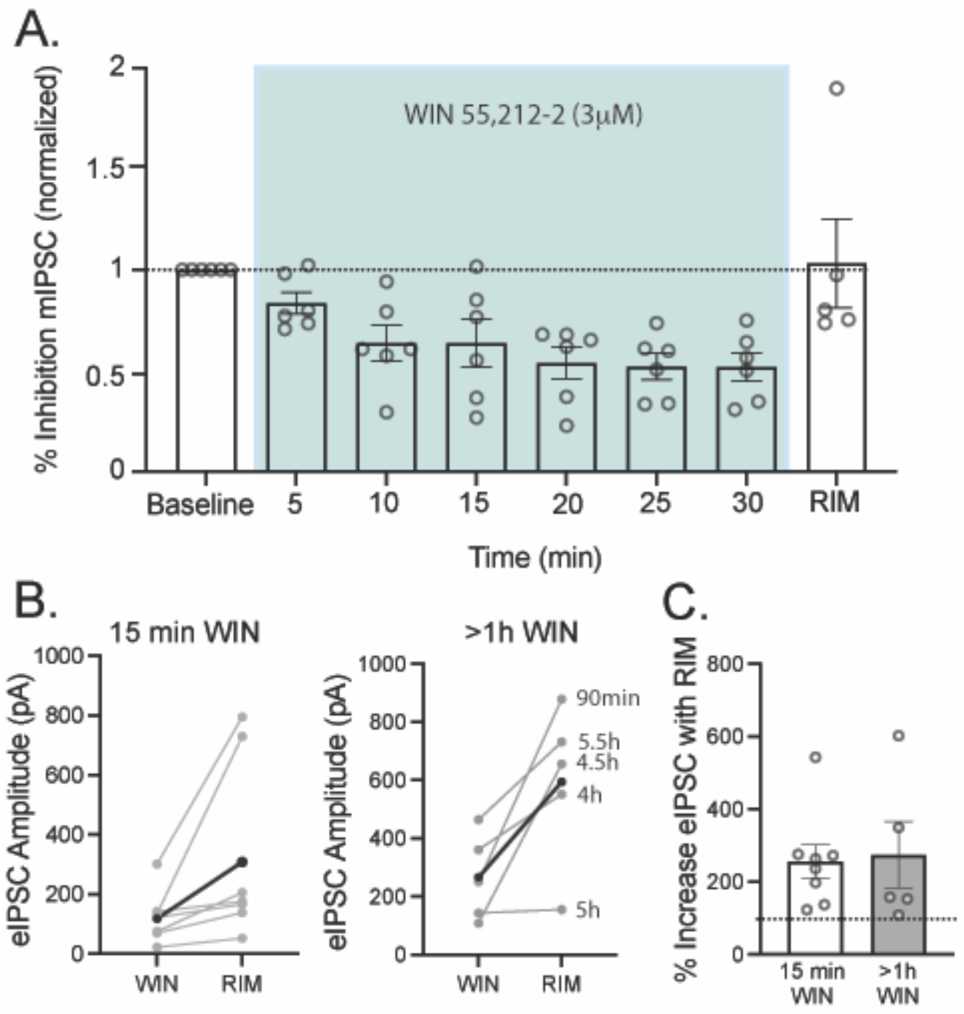
CB1R function is sustained throughout 5h WIN-induced activation (A) Percent inhibition of mIPSC frequency in vlPAG neurons during 30 min of WIN exposure (3 μM; n=8). Data are normalized to mIPSC frequency during baseline in TTX (500 nM) and NBQX (5 μM). WIN (3 μM) reduced mIPSC frequency over the first 10 minutes of drug application. Frequency remained reduced for the entirety of the 30 min drug application and was reversed by RIM (3 μM; two-tailed paired t-test, *t*_7_= 7; p=0.016). (B) eIPSC amplitude with bath application of CB1R selective antagonist RIM (3μM) after 15 minutes in WIN (3 μM; paired t-test, *t*_7_=2.42; p=0.046; data from 6 animals) or >1h WIN incubation (3 μM; paired t-test, *t*_5_=3.53; p=0.02; 5 cells from 3 animals). Average is shown in thick black. (C) Bar graph depicting RIM percent increase from WIN after 15 minutes in WIN (white bar) or >1 hour in WIN (gray bar; unpaired t-test, *t*_11_=0.2; p=0.8). Error bars represent SEM, dots indicate individual neurons.

### Persistent inflammation induces phosphorlylation dependent CB1R desensitization

Although CB1Rs are resistant to desensitization with acute agonist application over multiple hours (Fig. 4), it is possible that CB1R desensitization is induced by endogenous agonist(s) over the course of 5-7 days. A key step in canonical postsynaptic GPCR desensitization is G protein-coupled receptor kinase (GRK) phosphorylation of the GPCR C-terminal tail (Kovoor et al 1998, Lefkowitz 1993, Wang 2000, Zhang et al 1998). To block this step, we incubated slices in Compound 101 (Cmp101, 1 µM, ≥1 h), a potent and membrane permeable inhibitor of GRK 2/3 (Ikeda et al 2007, Thal et al 2011). Incubating slices in Cmp101 recovered CB1R suppression of GABA release after persistent inflammation (Fig. 5A; 41 ± 5% inhibition compared to CFA vehicle: 9 ± 2% inhibition). This result indicates that persistent inflammation induces GRK2/3-dependent desensitization of the presynaptic CB1R. We also tested Cmp101 (30 μM) incubation on CB1R function (Adhikary et al 2022, Leff et al 2020, Lowe et al 2015) and found that Cmp101 increased CB1R function in a concentration-dependent manner (30 μM incubation >1h, WIN inhibition 62 ± 10%). Interestingly, GRK2/3 desensitization after persistent inflammation appears to be selective to the CB1R as presynaptic MOR suppression of GABA release after Cmp101 incubation is not different between slices from either naïve or CFA-treated rats (30 μM; Fig. 5B).

**Figure 5:**
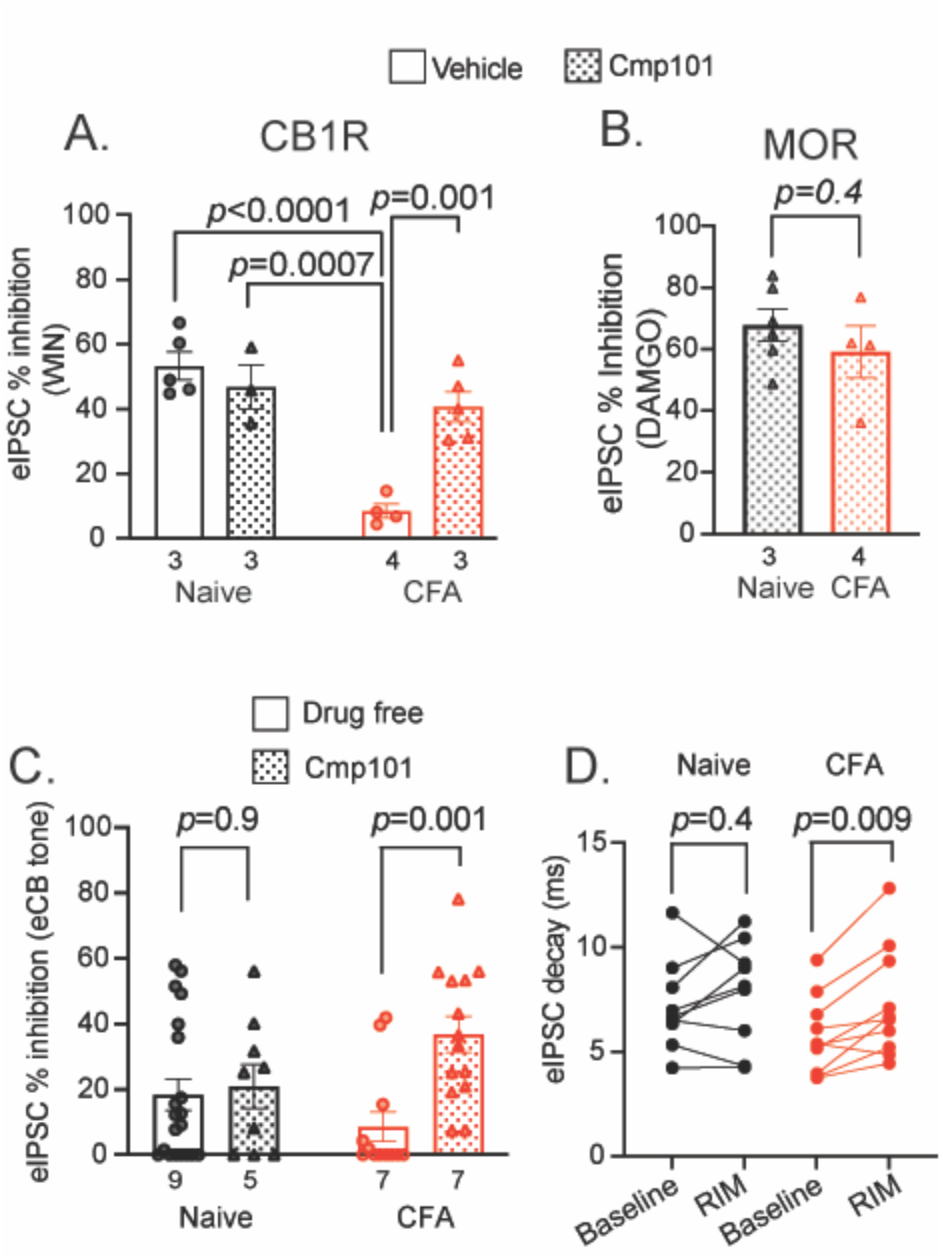
Compound 101 (Cmp101) incubation recovers CB1R inhibition of GABA release after persistent inflammation (A) WIN (3 μM) inhibition of eIPSC amplitudes from naïve (black) or CFA-treated (red) rats. vlPAG slices were incubated in vehicle (no fill) or Cmp101 (filled bar) for >1h. Cmp101 incubation fully recovered CB1R signaling in slices from CFA-treated rats (2-way ANOVA: main effect of Cmp101: F(1,13)=7.6; *p*=0.016; main effect of CFA: F(1,13)=29.9; *p*=0.0001; CFA x Cmp101 interaction: F(1,13)=17.29, p=0.001). Post-hoc analysis (Tukey test) reveals the effect of WIN in CFA-treated slices incubated in vehicle was significantly reduced compared to all other conditions. (B) DAMGO (1 μM) inhibition of eIPSC amplitude after Cmp101 (30 μM) incubation from naïve (black bar) or CFA-treated (red bar) rats. (C) Cmp101 incubation reveals eCB tone in recordings from CFA-treated rats (2-way ANOVA: main effect of Cmp101: F(1,46)=6.06; p=0.02). Post-hoc analysis (Šidák’s multiple comparisons test) reveals a significant effect of Cmp101 in CFA-treated rats but not naïve. RIM and NESS are combined. (D) eIPSC decay at baseline and after addition of RIM in slices from naïve (black) and CFA-treated (red) rats (2-way ANOVA: main effect of RIM F(1,17)=9.98; p=0.006; Šídák post-hoc test). Error bars represent SEM, dots indicate individual neurons and numbers represent the number of animals per bar.

The next experiments examined the role of eCB levels on the desensitization of CB1Rs after inflammation. GABAergic IPSCs were evoked and RIM (3 μM) was applied to evaluate tonic activation of CB1Rs by eCBs. Consistent with previous findings in the vlPAG (Aubrey et al 2017), RIM did not consistently increase eIPSC amplitude in recordings from slices from naïve rats (paired t-test: baseline vs. RIM: *t*_13_=1.54; *p*=0.15), nor was there consistent eCB tone in slices from CFA-treated rats (paired t-test: baseline vs. RIM: *t*_11_=1.13; *p*=0.28). Since inflammation induces CB1R desensitization, we hypothesized that eCB tone is masked by CB1R desensitization in rats treated with CFA. This was tested by incubating slices in Cmp101 prior to RIM superfusion. Cmp101 incubation did not reveal eCB tone in slices from naïve animals (Fig. 5C; naïve drug free: 13 ± 5% inhibition; naïve Cmp101: 12 ± 6% inhibition) but revealed significant eCB tone in slices from CFA-treated animals, (Fig. 5C; CFA drug free: 16 ± 6% inhibition; CFA Cmp101: 37 ± 6% inhibition). RIM is an inverse agonist, so we also tested eCB tone with the CB1R neutral antagonist, NESS 0327 (NESS; 0.5μM) to determine if the increased eCB tone resulted from constitutive activity of the CB1R (Ruiu et al 2003).

After Cmp101 incubation, NESS revealed eCB inhibition (40 ± 8%; n=9) which was similar to that produced by RIM (30 ± 5%; n=5). Thus, constitutive activity of CB1Rs does not account for the effect of the inverse agonist, RIM. A closer analysis of eIPSC kinetics revealed a decrease in eIPSC decay in recordings from CFA-treated rats, consistent with eCB modulation of vesicle release mode, changing multi-vesicular release to univesicular release in the vlPAG (Aubrey et al 2017). Even in the absence of Cmp101, RIM significantly increased eIPSC decay time in vlPAG slices from CFA-treated rats while it has no impact on decay in slices from naïve rats (Fig. 5D).

### Persistent inflammation prolongs 2-AG signaling in the vlPAG

With evidence that eCBs change eIPSC decay kinetics in the absence of Cmp101 (Fig. 5D), it appeared that eCBs activate CB1Rs even though the majority are desensitized after persistent inflammation (Fig. 5A). We used depolarization-induced suppression of inhibition (DSI) to directly examine eCB activation of CB1Rs. The DSI protocol (+20mV for 5 seconds; (Wamsteeker et al 2010)) induced a rapid and transient suppression of presynaptic GABA release in a subset of PAG neurons (Fig. 6A). This suppression was blocked by the CB1R antagonist NESS (0.5 μM; Fig. 6A), indicating that DSI induces CB1R activation in the vlPAG. In slices from CFA-treated rats, we observed prolonged DSI (Fig. 6B) that was also blocked by NESS (Fig. 6B). The prolonged time course was analyzed by measuring the maximal percent inhibition immediately following depolarization (max DSI) and 30 s later (late DSI; Fig. 6C). Max DSI in recordings from naïve rats is 41 ± 6% is similar to 37 ± 4% in recordings from CFA-treated rats. The eIPSCs from naïve slices return close to baseline (14 ± 5% inhibition) but did not recover in 30 s (37 ± 5%) in recordings from CFA treated rats (Fig. 6C). In addition to prolonged DSI, the proportion of experiments that yield DSI after depolarization is significantly increased (Fig. 6D) in recordings from CFA treated rats (16 out of 19 cells exhibited DSI) compared to recordings from naïve rats (10 out of 24 cells exhibited DSI). In the remaining neurons, the DSI protocol had no effect on eIPSC amplitude (Fig. S3A).

**Figure 6:**
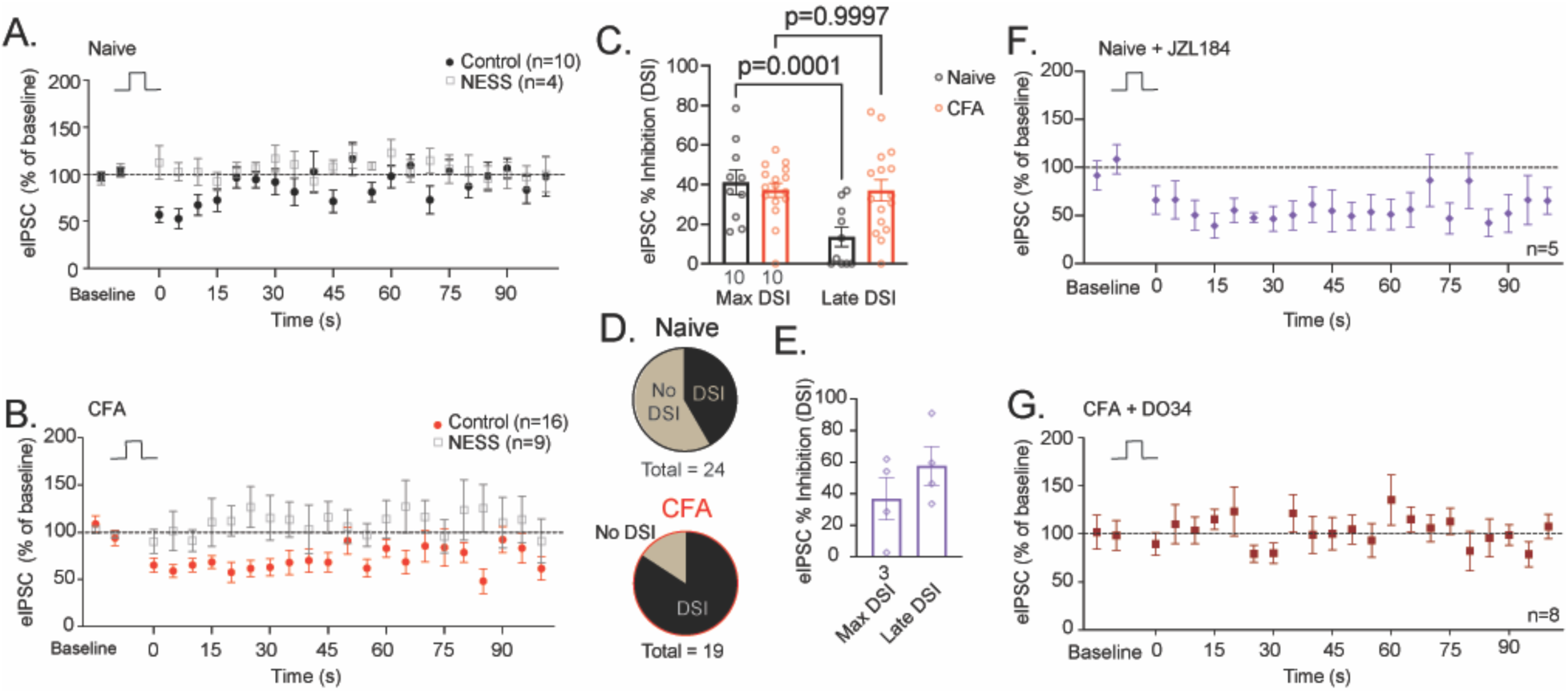
Persistent inflammation increases 2-AG activity at the CB1R. (A) Summary of DSI (5s; +20 mV) in tissue from naïve rats (black circles; n=10 recordings from 10 rats). DSI is blocked by NESS (0.5μM; gray open boxes; n=4 recordings from 3 rats) (B) Summary of DSI in tissue from CFA-treated rats (red dots; n=16 recordings from 10 rats). DSI is blocked by NESS (0.5μM; gray open boxes; n=9 recordings from 5 rats).(C) Quantification of eIPSC % inhibition at max DSI and late DSI in vlPAG tissue from naïve (black) and CFA-treated (red) animals (2-way repeated-measures ANOVA: main effect DSI length: F(1,24)=14.5; p=0.0009; interaction DSI length x CFA: F(1,24)=14.3; p=0.0009; Šídák post-hoc test). (D) Proportion of patched neurons that respond to DSI. In slices from naïve rats, after depolarization 10 neurons exhibited DSI and 14 did not. In slices from CFA treated rats, 16 exhibited DSI and 3 did not. The proportion of neurons that produced DSI was significantly higher in slices from CFA-treated slices (Fishers exact test: p=0.006). (E) Quantification of eIPSC % inhibition at max DSI and late DSI in vlPAG tissue from naïve animals incubated in MAGL inhibitor JZL184 (1 μM, >1h). (F) Summary of DSI in tissue from naïve rats after incubation in JZL184 (1 μM, >1h). (G) Summary of DSI in tissue from CFA-treated rats incubated in DAGLα inhibitor, DO34 (1 μM incubation; >1h). Dots represent individual recordings, numbers below the bar represent number of animals; error bars represent SEM.

DSI is dependent on 2-AG signaling in other brain regions and can be prolonged by inhibiting 2-AG degradation (Hashimotodani et al 2007, Straiker & Mackie 2005). To determine the effects of prolonging 2-AG levels in vlPAG, slices from naïve rats were incubated in the 2-AG degradation inhibitor, JZL184 (1μM) for at least one hour (Lau et al 2014, Long et al 2009)). Incubation with JZL184 prolonged DSI and recapitulated the DSI time course observed in recordings from CFA-treated rats (Fig. 6E,F). Interestingly, JZL184 incubation did not change the proportion of cells that exhibit DSI (Fig. S3B). DSI after CFA is completely blocked by inhibiting 2-AG synthesis with incubation in the DAGLα inhibitor, DO34 (Fig. 6G; 1μM; >1h). Together, these data indicate that CFA-induced inflammation increases 2-AG levels in the vlPAG.

## Discussion

Here, we describe a mechanism by which persistent inflammation induces adaptations in the endogenous cannabinoid system. Inflammation promotes desensitization of presynaptic CB1Rs that suppress GABA release in the vlPAG. This desensitization is dependent on CB1R and GRK2/3 activity and is recovered in the presence of the GRK2/3 inhibitor, Cmp101. Cmp101 also reveals an underlying increase in tonic activation of CB1Rs by eCBs 5-7 days after CFA injection. Despite these adaptations, desensitization does not affect maximal CB1R activation by eCBs, but actually prolongs CB1R activation by depolarization-induced eCB release in CFA-treated rats. These data have important implications for the development of pharmaceuticals targeting the cannabinoid system for inflammatory diseases.

Our data show direct evidence of GRK2/3-dependent desensitization of presynaptic CB1Rs. While postsynaptic GPCRs readily desensitize and internalize in response to agonist exposure (Williams et al 2013), it is well established that presynaptic GPCRs are resistant to desensitization (Fyfe et al 2010, Pennock et al 2012, Pennock & Hentges 2011, Pennock & Hentges 2016, Wetherington & Lambert 2002). Sustained signaling from presynaptic receptors during prolonged agonist exposure may be due to multiple mechanisms. One such mechanism involves protein-protein interactions with presynaptic scaffold proteins that immobilize the receptors close to the plasma membrane, as observed for presynaptic GABA_B_ receptors (Boudin et al 2000, Laviv et al 2011, Vargas et al 2008, Vigot et al 2006). An alternative mechanism, observed for presynaptic MORs, describes presynaptic GPCRs internalizing into endosomes in response to agonist stimulation, but maintaining signaling through rapid receptor replacement by a pool of receptors that diffuse laterally through axon membranes (Jullie et al 2020). Both mechanisms result in sustained GPCR signaling from presynaptic terminals. CB1Rs exhibit rapid mobility through the synapse under basal conditions; however, in contrast to MOR regulation, prolonged agonist exposure significantly reduces CB1R mobility and expression of CB1Rs in the synapse (Mikasova et al 2008). We demonstrated that CB1Rs are also resistant to desensitization under normal conditions but are desensitized after persistent inflammation, in contrast to presynaptic MORs which were unaffected by inflammation. Differences in CB1R and MOR regulation and mobility could underly their differential desensitization after persistent inflammation.

CB1R desensitization in response to prolonged administration of exogenous agonists, such as tetrahydrocannabinol (Δ^9^-THC) or WIN, has been reported by many groups (Breivogel et al 1999, Kouznetsova et al 2002, Lazenka et al 2014, Rubino et al 2000, Sim et al 1996). Long-term increases in eCBs also induce CB1R desensitization (Imperatore et al 2015, Kinsey et al 2013, Long et al 2009, Navia-Paldanius et al 2015, Schlosburg et al 2010). eCB levels in the PAG are increased almost immediately after acute inflammation induced by formalin injection into the hindpaw (Walker et al 1999) as well as after 7 days of chronic constriction injury, a model of neuropathic pain (Petrosino et al 2007). The observed CB1R desensitization in our study is likely a result of increased CB1R-induced G protein signaling within the vlPAG early in inflammation (Wilson-Poe et al 2021).

We observed prolonged DSI after persistent inflammation, which is consistent with the time course in other studies pharmacologically or genetically inhibiting MAGL (Chen et al 2016, Pan et al 2009, Schlosburg et al 2010, Straiker & Mackie 2005). We show that the prolonged inhibition of GABAergic IPSCs following DSI in slices from CFA-treated rats is blocked by inhibiting DAGLα, the enzyme responsible for 2-AG synthesis, implicating 2-AG in the adaptations induced by CFA in the vlPAG. The prolonged time course could be the result of increased synthesis or decreased activity or levels of the degradation enzyme, MAGL. Since we observe a comparable maximal effect of DSI in recordings from both naïve and CFA-treated rats, suggesting comparable levels of 2-AG synthesis, we hypothesize that MAGL activity is diminished following persistent inflammation. Under normal conditions in the vlPAG, MAGL catabolizes 2-AG quickly enough that washing 2-AG over the slice does not suppress GABA release unless MAGL is blocked (Lau et al 2014). Consistent with this interpretation, experiments using MAGL knockout mice or pharmacological inhibition of MAGL show increases in 2-AG signaling that lead to CB1R desensitization (Imperatore et al 2015, Kinsey et al 2013, Long et al 2009, Navia-Paldanius et al 2015, Schlosburg et al 2010). However, if alterations in MAGL degradation of 2-AG are the sole mechanism underlying these adaptations in CFA-treated rats, we expected inhibiting MAGL activity with JZL184 would also increase the proportion of neurons in naïve rats that displayed DSI. This was not the case suggesting that CFA treatment may also affect synthesis in neurons that do not readily display DSI or diffusion of eCBs within the vlPAG. Therefore, reductions in MAGL degradation of 2-AG play a role but other mechanisms may be also involved in inflammation-induced adaptations in the cannabinoid system.

These results also highlight differences in signaling between exogenous and endogenous cannabinoids following persistent inflammation. Desensitization of CB1Rs clearly diminishes effects of exogenous cannabinoid agonists but eCBs continue to activate CB1Rs and induce prolonged suppression of GABA release, even though the majority of CB1Rs are desensitized. Similar reductions in exogenous but not endogenous ligand-mediated CB1R suppression of GABA release have been observed after chronic stress paradigms (Patel et al 2009). Importantly, this indicates that eCBs synthesized through DSI protocols are coupled more effectively to effectors and may indicate spare receptors in synapses. Alternatively, eCBs target different signaling pathways. Further studies are necessary to understand the consequences of long-term alterations in eCB synthesis and CB1R desensitization.

### Physiological Relevance

Direct microinjections of cannabinoid agonists into the PAG induce antinociception (Lichtman et al 1996, Martin et al 1995, Wilson et al 2008, Wilson-Poe et al 2013) through activation of CB1Rs that inhibit GABA release in the PAG (Vaughan et al 2000). Recent work has highlighted MAGL inhibitors as analgesic therapeutic options (Anderson et al 2014, Curry et al 2018, Della Pietra et al 2021, Diester et al 2021, Ignatowska-Jankowska et al 2015) but the data presented here suggest that MAGL inhibition may not be a viable strategy if inflammation impairs MAGL function and desensitizes CB1Rs. However, systemic administration of MAGL inhibitors, as well as fatty acid hydrolase (FAAH) inhibitors and combinations of the two, increase levels of the eCBs 2-AG and anadamide and result in anti-hyperalgesia in both neuropathic and inflammatory pain models (Anderson et al 2014, Jayamanne et al 2006, Mitchell et al 2005). In addition, the anti-hyperalgesic effects of systemic cannabinoid agonist, Δ^9^- THC (Craft et al 2013, Smith et al 1998, Sofia et al 1973), and WIN (Bridges et al 2001, Herzberg et al 1997, Li et al 1999) are not altered in similar inflammatory or neuropathic pain models, suggesting either that the reduced functional CB1Rs in the vlPAG are sufficient for cannabinoid-induced analgesia or that CB1Rs in the vlPAG are not required. One intriguing possibility is that inflammation-induced increases in 2-AG contribute to hyperalgesia and CB1R desensitization is a compensatory response that protects synapses. Indeed, there is precedent for cannabinoids to contribute to hyperalgesia (Dunford et al 2021, Khasabova et al 2022). Understanding the behavioral consequences of this altered cannabinoid signaling within the vlPAG after persistent inflammation, the generalizability to other brain areas, and the reversibility of this process have important implications for future drug development.

## Materials and Methods

### Animals

Adult male and female Sprague Dawley rats (Harlan Laboratories and bred in-house; postnatal day 30-90) were used for all experiments. All procedures were performed in strict accordance with the *Guide for the Care and Use of Laboratory Animals* as adopted by the Institutional Animal Care and Use Committee of Oregon Health & Science University. Care was taken to minimize discomfort.

### Inflammation

Complete Freund’s Adjuvant (CFA: heat-killed *Mycobacterium tuberculosis* in mineral oil, 1 mg/ml, 0.1 ml volume injected, Sigma-Aldrich) was injected subcutaneously into the plantar surface of the right hindpaw. The CFA injection produces an intense tissue inflammation of the hindpaw characterized by erythema, edema, and hyperalgesia (Iadarola et al 1988). Electrophysiological recordings and tissue dissections were performed 5-7d following CFA injection.

### vlPAG slice preparation

vlPAG slices were prepared as previously described (Tonsfeldt et al 2016). Rats were deeply anesthetized with isofluorane and the brain was rapidly removed and placed in ice-cold sucrose-based cutting buffer containing the following (in mM): 75 NaCl, 2.5 KCl, 0.1 CaCl_2_, 6 MgSO_4_, 1.2 NaH_2_PO_4_, 25 NaHCO_3_, 2.5 dextrose, 80 sucrose. Ventrolateral PAG (vlPAG) slices were cut to a thickness of 220 µm on a vibrotome (Leica Microsystems) in sucrose-based cutting buffer and transferred to a holding chamber with aCSF containing the following (in mM): 126 NaCl, 21.4 NaHCO_3_, 22 dextrose, 2.5 KCl, 2.4 CaCl_2_, 1.2 MgCl_2_, 1.2 NaH_2_PO_4_ and the osmolarity was adjusted to 300-310 mOsm. Slices were maintained with 95% O_2_- and 5% CO_2_-oxygenated until transfer to a recording chamber on an Olympus BX51WI upright microscope and superfused with aCSF maintained at 32°C.

### Whole-cell patch-clamp recordings

Voltage-clamp recordings (holding potential -70 mV) were made in whole-cell configuration using an Axopatch 200B amplifier (Molecular Devices). Patch-clamp electrodes were pulled from boroxilicate glass (1.5 mm diameter; WPI) on a two-stage puller (PP83, Narishige). Pipettes had a resistance of 2.5-5 MΩ. IPSCs were recorded in an intracellular pipette solution containing the following (in mM): 140 CsCl, 10 HEPES, 4 MgATP, 3 NaGTP, 1 EGTA, 1 MgCl_2_, 0.3 CaCl_2_, pH adjusted to 7.3 with CsOH, 290-300 mOsm. QX314 (100μM) was added to the internal solution for eIPSC experiments to reduce action potentials in the recording cell. Access resistance was continuously monitored. Recordings in which access resistance changed by >20% during the experiment were excluded from data analysis. A junction potential of -5mV was corrected during recording. GABAergic events were isolated in the presence of glutamate receptor antagonist NBQX (5 µM). Spontaneous miniature IPSCs (mIPSCs) were recorded in the presence of TTX (500 nM). Events were low-pass filtered at 2 kHz and sampled at 10-20 kHz for off-line analysis (Axograph 1.7.6) and individual events were visually confirmed. In experiments using exogenous cannabinoid agnoists, one neuron was recorded per slice due to the lipophilic nature of cannabinoid receptor drugs. After each experiment with exogenous cannabinoid agonists or antagonists, lines were washed with 50% EtOH. Each set of experiments was repeated using at least 3 distinct rats with no more than 2 cells from a single rat included in a specific dataset.

### Drugs

WIN55,212-2 (Caymen Chemicals), SR141716A (rimonabant; RIM; Caymen Chemical), and NESS (Tocris) were dissolved in DMSO, aliquoted, and stored at -20°C. CP55,940 and AM251 (Caymen Chemical Company) were dissolved in methanol and stored at - 20°C. DMSO and methanol at appropriate concentrations were used as vehicle controls. 2,3-dihydroxy-6-nitro-7-sulphamoyl-benzo(F)quinoxaline (NBQX; (Sheardown et al 1990)), [D-Ala^2^, N-MePhe^4^, Gly-ol]-enkephalin (DAMGO), Naloxone and tetrodotoxin (TTX) were purchased from Abcam, dissolved in distilled water, and stored at 4°C. CMP101 (3- [(4-methyl-5- pyridin-4-yl-1,2,4-triazol-3-yl)methylamino]-N-[[2- (trifluoromethyl) phenyl]methyl]benzamide hydrochloride) was purchased from Hello Bio and prepared as described previously (Leff et al 2020). Briefly, Cmp101 (made fresh daily) was first dissolved in a small amount of DMSO (10% of final volume), sonicated, then brought to its final volume with 20% (2-Hydroxypropyl)-b-cyclo-dextrin (HPCD; Sigma-Aldrich) and sonicated again to create a 10mM solution. For experiments using a higher concentration of Cmp101, Cmp101 was applied to the slice as follows: 30μM incubation for >1h, 1μM maintenance while patching, 10μM in drug tubes (Adhikary et al 2022, Leff et al 2020, Lowe et al 2015). For experiments using a lower concentration of Cmp101, [1μM] was used for incubation (>1h), maintenance while patching, and in drug tubes. DMSO and 20% HPCD were used as the vehicle control.

### Radioligand Binding Assay – tissue dissection

Rats were deeply anesthetized with isoflurane, brains were removed and submerged in ice cold Tris-HCl buffer (pH=7.4 at 4°C). Over ice, the brain was sectioned into 1mm slices from which the vlPAG, DLS and hypothalamus were dissected and immediately flash frozen on dry ice. Tissue samples were stored at -80°C.

### Radioligand Binding Assay- total particulate tissue preparation

Tissue preparation was adapted from (Eastwood et al 2018). Since brain regions sampled are so small, tissue from each brain region from multiple animals (8 vlPAG, 2 DLS, 2 hypothalamus) was pooled to ensure ample protein levels for saturation binding. Tissue was removed from -80°C and transferred to 2 ml tube containing 0.5 ml Tris-HCl (pH 7.4 at 4C) with protease inhibitor (EMD Millipore; protease inhibitor cocktail set #539134). Tissue was homogenized with a polytron PT1200E 4 x 6s, placing sample on ice for 20s between homogenizations. The polytron was washed with water between each sample. The volume was increased to 1.5ml, then the sample was transferred to a mini-centrifuge and spun at 13,000 x g for 20 min at 4°C. The supernatant was discarded and pellet was resuspended in 0.5ml Tris-HCl with protease inhibitor. Tissue was homogenized for 7s and spun as described above once more. After the final spin, the supernatant was discarded, the pellet was resuspended in TME Binding Buffer (200 mM Tris Base, 50mM MgCl_2_, 10mM EDTA, pH=8.0) with protease inhibitor and homogenized for 10s. TME with protease inhibitor was added for a final volume of 1.5ml. Samples were kept on ice throughout the preparation. Protein levels were determined with the BCA Protein Assay Kit (Thermo Fisher Scientific, Waltham, MA).

### Radioligand Binding Assay- Saturation Curve

Binding assays were conducted in the absence of Na^+^. [^3^H]CP-55,940 was used to measure cannabinoid receptor binding (Catani & Gasperi 2016, Chanda et al 2010, Freels et al 2020, Hill et al 2008, McLaughlin et al 2013, Romero et al 1995). Binding assays were conducted using 5 concentrations [^3^H]CP-55,940 (0.1-7nM) in a final volume of 1 ml. Assays were performed in duplicate in a 96-well plate with 50 mM TME binding buffer with bovine serum albumin (BSA; 1mg/ml; pH 7.4 at 30°C). Nonspecific binding was subtracted from total binding to yield specific binding. Nonspecific binding was determined with 1μM WIN55,212-2 and was 59%, 19%, or 55% in naïve and 53%, 17%, or 55% in CFA in vlPAG, DLS, and hypothalamus, respectively. Prepared membranes were incubated with [^3^H]CP-55,940 at 30°C for 60 min. The incubation was terminated using a Tomtec cell harvester (Hamden, CT) by rapid filtration through Perkin Elmer Filtermat A filters presoaked in 0.2% polyethylenimine. The filters were dried, spotted with scintillation cocktail, and radioactivity was determined using a Perkin Elmer microBetaplate 1405 scintillation counter.

### Data Analysis

In all electrophysiological experiments, each dataset included recordings from at least 3 rats. For DSI experiments, “Max DSI” averaged the first 4 eIPSCs after depolarization and “Late DSI” averaged eIPSCs 30-45 seconds after depolarization. In radioligand binding experiments, 3 replicates per group were run. All analysis were conducted in Graphpad Prism 9 (Prism version 9.2; San Diego, CA). Values are presented as mean ± SE and all data points are shown in bar graphs to illustrate variability. Statistical comparisons were made using two-tailed paired or unpaired T-test, one-way ANOVA, or two-way ANOVA when appropriate. In all summary bar graphs, each dot represents an individual cell while the numbers in the bars represent the animal number. When post-hoc analysis was appropriate Tukey test and Šidák’s multiple comparisons tests were used. P < 0.05 was used.

## Acknowledgments

We would like to thank Dr. Amy Eshleman for her expertise and guidance with radioligand binding assays. We also thank Dr. John Williams and Dr. Sweta Adhikary for help with the Cmp101 experiments. We thank members of the Ingram laboratory, as well as Dr. Mary Heinricher and her laboratory, for valuable discussion and suggestions. Work was supported by funding from NIH R01DA042565 (S.L.I.), C.A.B. supported by research grants from the NIH/NIDA (F31 DA052114 and T32 DA007262). This work was supported by funding to AJ from NIH/NIDA (ADA20003-001-00002), from the US Drug Enforcement Administration (D-20-OD-00), from the US Food and Drug Administration (CDER-20-I-0546), and from the Department of Veterans Affairs Research Career Scientist Program (1IK6BX005754).

## Author Contributions

C.A.B. performed experiments and C.A.B. and S.L.I. conceived of the experiments, analyzed the data and wrote the manuscript. A.J. provided essential reagents, equipment, and helped with analysis of radioligand binding assays.

