## Supplemental Figures for "Persistent inflammation promotes endocannabinoid release and presynaptic cannabinoid 1 receptor desensitization"

**Supplemental Figure 1**

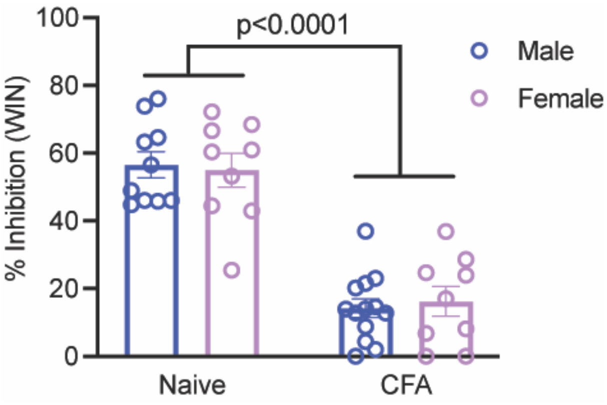


Figure S1: No sex differences in WIN suppression of GABA release in naïve or CFA-treated animals. WIN inhibition of all eIPSCs (Fig. 1), mIPSCs (Fig. 2), and vehicle eIPSCs (Fig. 5) were pooled and separated by sex. 2-way ANOVA reveals a significant main effect of CFA ($F(1,37)=107$; $p<0.0001$) and no sex difference ($p=0.96$). Dots represent individual recordings, error bars represent SEM.

**Supplemental Figure 2**

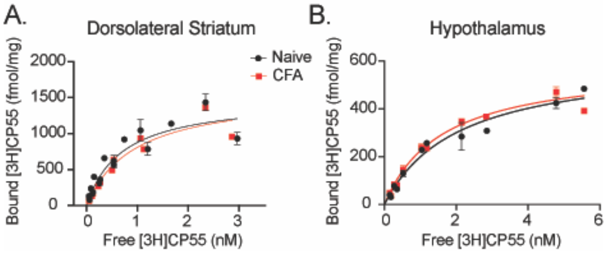


Figure S2- Cannabinoid receptor binding is not affected by persistent inflammation in multiple brain regions. [$^3$H]CP-55,940 binding in the (A) dorsolateral striatum and (B) hypothalamus dissected from brains of naïve (black) and CFA-treated (red) rats.

### Supplemental Figure 3

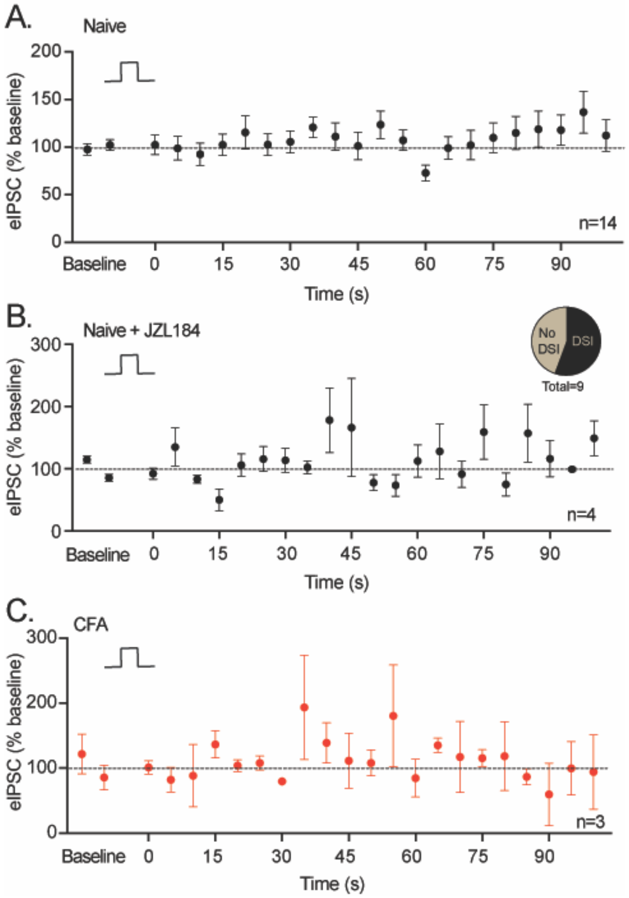


Figure S3- Summary of trace from naïve and CFA-treated slices where depolarization (5s; +20mV) did not induce suppression of inhibition. (A) Naïve; (B) Naïve incubated in JZL184 (1μM; >1h); out of 9 recordings 5 had DSI and 4 did not (inset). (C) CFA-treated animal. Error bars represent SEM.
